# Microbiome Assembly in *Zostera marina* during early host development: An EcoFAB 2.0 Application for Aquatic Plant-Microbe Interactions

**DOI:** 10.1101/2025.08.27.672729

**Authors:** Gina Chaput, Emma A. Deen, Eric Q. Pham, Sarah Crystal, Andrew Matayoshi, Peter Andeer, Trent R. Northen, Jonathan A. Eisen, John J. Stachowicz, E. Maggie Sogin

## Abstract

Seagrass restoration practices are evolving to leverage microbiome applications, similar to agricultural systems that have demonstrated how targeted microbial communities enhance crop resilience in challenging environments. While adult seagrass microbiome research has expanded significantly, research on the seed microbiome remains critically understudied. This gap is particularly important given that seeds represent a large portion of restoration efforts. Advancing seed microbiome research requires standardized experimental systems for controlled plant-microbe interaction studies, which are currently lacking in seagrass research. Here, we tested fabricated ecosystem devices (EcoFAB 2.0) as a standardized system for growing seedlings of *Zostera marina* (eelgrass), enabling a controlled study of aquatic plant-microbe interactions. Using these chambers, we addressed three key questions: (i) Can we reliably grow eelgrass in a controlled laboratory setting? (ii) Can we manipulate eelgrass microbiome assembly and its long-term trajectory? and (iii) Can we detect shifts in the microbiome during plant development (host filtering)? Host morphology measurements and 16S rRNA gene amplicon sequencing were used to track microbiome assembly across three early developmental stages of the host. We identified 143 ASVs that differed between plants from surface-sterilized versus intact seeds, highlighting the strong influence of epiphytic bacteria on microbiome assembly. We also identified 26 stage-specific indicator ASVs across eelgrass development, suggesting stage-specific microbial associations during seedling establishment. This work demonstrates the potential for targeted manipulation of the microbiome in seagrass for restoration efforts.

**Importance:** Using the Fabricated Ecosystem 2.0 (EcoFAB 2.0), we were able to successfully control the microbial environment of *Z. marina*, resulting in the removal of epiphytes and maintaining low microbial diversity across plants without compromising the growth dynamics of the seedlings. Our findings advance the marine plant model system, *Zostera marina*, by identifying taxonomic indicators across life stages. This work lays the foundation for a targeted understanding and application of microbiomes for seagrass restoration, bridging the critical knowledge gap between agricultural seed microbiome success and marine restoration applications.

## Introduction

Seagrasses, including *Zostera marina* (also known as eelgrass), are foundation species that occur along coastlines and serve as nurseries for commercial and ecologically important organisms, protect coastlines from erosion, and drive biogeochemical cycling of nutrients^1,2^. Unfortunately, anthropogenic activities are driving declines in seagrass meadows, which jeopardize the services they provide to coastal communities^3^. Efforts to restore deteriorated meadows or establish new habitats have had mixed success, with research reporting varying outcomes across different approaches and locations^4,5^. To achieve meaningful progress, it is critical to develop solutions that protect current seagrass populations, work alongside local communities to apply methods that work for both seagrass and local economies, and develop solutions that “future-proof” seagrass populations^6^. The resilience of seagrass beds could be enhanced through a range of approaches, including selective breeding, acclimatization, and food web manipulations. Additionally, the possibility of leveraging the microbiome to promote plant resilience and resistance to stressors has been proposed^6,7^. Terrestrial agroecosystems have a long history of application of microbiome science to increase crop yields, and these approaches have recently been piloted in some marine systems as well (e.g., corals, mangroves^8,9^), suggesting that microbes have untapped potential in seagrass restoration and conservation.

Seed microbiomes of crops, such as wheat, maize, and tomatoes, provide protection, promote germination, and aid in nutrient exchange with the environment as the seed begins to develop^10–12^. The growing recognition of seed microbiomes’ importance in plant success has led to the development of synthetic microbial communities that can be inoculated onto seeds for use in agriculture^13,14^. However, much less is known about the role of microbial species in promoting seagrass growth or seed development. Current understanding of the *Z. marina* microbiomes is largely based on adult plants^15–22^. It is unclear how the microbiome assembles on seagrass seeds and the functional roles of individual members once established^16,23^. If we want to apply microbiome treatments to help expand and restore seagrass meadows, we must first understand the assembly and roles of different microbial taxa in seed-associated processes. This includes determining their impacts on germination success and seedling survival.

To study seagrass seed microbiome assembly dynamics, it is necessary to develop sterile, fabricated system approaches that enable precise manipulation of the plant microbiome during seagrass development. Multiple mesocosm chamber designs have been developed for studying terrestrial plant-microbe interactions in the rhizosphere (see Yee *et al*. 2021 for a comprehensive review)^24–26^. Similarly, phyllosphere research has adopted various platforms alongside traditional approaches, including petri dishes, tissue culture boxes, and FlowPots/GnotoPots^27^. In contrast, seagrass research has relied on a limited suite of culture systems: petri dishes for seed germination^28^, glass bottles for seedling cultivation^29^, and flow-through tanks or two-compartment hydroponic chambers for adult plant growth^15,30^. Here, we adapt the Fabricated Ecosystem 2.0 (EcoFAB), a self-contained and dual-chamber platform^25^, to grow *Z. marina* in a fully submerged environment while tracking microbiome assembly throughout plant development. We were able to successfully control the microbial environment of *Z. marina*, resulting in the removal of epiphytic microbiomes and maintaining consistently low microbial diversity across plants without compromising the growth dynamics of the seedlings. These chambers allowed us to address the following questions: (i) Can we reliably grow eelgrass in a controlled laboratory setting, (ii) Can we manipulate the eelgrass microbiome assembly and its long-term trajectory successfully, and, (iii) Can we detect shifts in the microbiome during plant development (i.e., host filtering)?

## Methods and Materials

### Seed collection and treatment

*Z. marina* flowering shoots were collected from Bodega Bay, CA (38°20’00.6”N 123°03’34.7”W), in June, July, and August of 2023. To allow developing seeds to mature while preventing sediment-associated microbiome contamination, flowering shoots with intact rhizomes were rinsed and placed in 500 μm nylon mesh bags, then held in flow-through tanks at the UC Davis Coastal and Marine Sciences Institute’s Bodega Marine Laboratory until September. Seeds were sorted, and any with visible damage, pathogen growth (e.g., phytophthora), or softness under pressure were removed. The resulting 1,017 seeds were divided into a reduced microbiome (n=500) and intact microbiome (n=517) treatment group. To reduce the epiphytic microbiome, we adapted sterilization techniques used for terrestrial seed microbiome studies^13,31^: seeds were surface sterilized using a single wash each of 95% ethanol (15 seconds), 5% bleach (3 minutes), and 70% ethanol (2 minutes), followed by three washes with filter-sterilized (0.22 µm) Instant Ocean (32 PPT; 30 seconds each). Seeds with intact microbiomes were also shaken with filter-sterilized Instant Ocean (30 seconds each) to mimic scarification that the surface sterilization procedure may have caused^32,33^. To confirm bacterial load differences between the treatment groups, we tested our methods before the experiment by plating the third wash of Instant Ocean from each treatment group onto Marine Broth Agar and found that the reduced microbiome group had no colony-forming units, whereas the intact group did.

The reduced and intact seeds were allowed to either germinate in the dark or were under a 12 hr:12 hr light:dark cycle. All seeds prior to germinating were kept at 15 °C and placed into petri dishes filled with 8 mL of filter-sterilized (0.22 µm) Instant Ocean (32 PPT) and 5 mL of hydrogel beads (14G needle) ^34^. Seeds were checked every 3-4 days for initial germination, pathogen or algal overgrowth, and mortality. Germinated seeds were then randomly assigned to sterile petri dishes, test tubes, or EcoFAB 2.0 chambers^25^ (**Table S1**). Chambers were filled with filter-sterilized (0.22 µm) Instant Ocean (32 PPT), and the petri dishes, test tubes, and lower chamber of the EcoFab were filled with ∼5 mL of hydrogel beads to mimic sediments^34^. To keep both chambers of the EcoFABs water-tight for seagrass, a silicone gasket was cut and placed between the EcoFAB chamber and the EcoFAB base. All chambers were kept at 15 °C and maintained on a 12 hr:12 hr light:dark cycle. To avoid an oxygen gradient forming in the smaller volume petri dishes and test tubes, chambers were gently shaken for 30 minutes per day at 180 RPM.

### Eelgrass seed and seedling sampling and measurements

Across treatments, seeds and seedlings were randomly selected to be destructively harvested before introduction to the chambers (Stage 1: initial germination), at Stage 3 (first true leaf develops), or Stage 6 (cotyledon senesces and abscises, roots are formed, and at least two true leaves have appeared; **Table S1**) of development. For each seedling, each plant was photographed using a dissecting microscope and the cotyledon (cotyledonary blade, cotyledonary sheath, and axial hypocotyls), leaf, and root lengths were measured using ImageJ^35^. Following imaging, samples (entire plants) were stored at -80 °C until processed for DNA extractions. DNA was extracted using the DNeasy Plant Mini Kit (Qiagen, Hilden, Germany) following the manufacturer’s guidelines.

### 16S rRNA gene library construction and sequencing

DNA was prepared for V3/V4 amplification of 16S rRNA genes using the Zymo Quick-16S Plus NGS Library Prep Kit and unique dual indexes (Zymo Research, CA, USA). Primers used by the Zymo Quick-16S Plus NGS Library Prep Kit are 341F (CCTACGGGDGGCWGCAG, CCTAYGGGGYGCWGCAG; 17 bp) and 806R (GACTACNVGGGTMTCTAATCC; 21 bp). PCR conditions were: 95 °C for 10 minutes, 35 cycles of [95 °C for 30 seconds, 55 °C for 30 seconds, 72 °C for 3 minutes], and 72 °C for 6 minutes. Sequencing was performed on the Illumina NextSeq2000 platform using a 600-cycle flow cell kit to produce 2 x 300 bp paired reads (SeqCoast Genomics, NH, USA). A 30-40% PhiX control (unindexed) was spiked into the library pool to support optimal base calling of low diversity libraries on patterned flow cells. Read demultiplexing, read trimming, and run analytics were performed using DRAGEN v3.10.12^36^, an onboard analysis software on the NextSeq2000.

### Data analyses for seedling morphology measurements and sequence data

A general linearized model was used to determine how seed (i.e., reduced vs. intact microbiome) and light treatments (i.e., dark or a 12 hr:12 hr light:dark cycle) affected seed germination success. To compare variation in plant growth (cotyledon, leaf, and root length) as a function of seed treatment and chamber type (i.e., test tube, petri dish and EcoFAB), a combination of parametric (i.e., 2-way ANOVA, mixed linear effects model) and non-parametric approaches (i.e., Scheirer–Ray–Hare test) was used. For Stage 3 leaf lengths, a linear mixed effects model (LMM fit by REML) that is robust to heteroscedasticity was applied, while for Stage 3 cotyledon lengths, a Scheirer–Ray–Hare test followed by a Dunn’s post hoc test was used. Stage 6 leaf measurements were tested with a 2-way ANOVA, and root length was tested with Scheirer–Ray–Hare test followed by a Dunn’s post hoc test. Effects of seed treatment and chamber type on days until the plant developed to either Stage 3 or 6 were tested with Scheirer–Ray–Hare test, followed by a Dunn’s post hoc test. All statistics described above were analyzed in R Statistical Software (v.4.2.1)^37^ and RStudio^38^ (v.2022.7.1.554) with the vegan (v.2.6-4)^39^, rcompanion (v.2.3.0)^40^, and dunn.test (v.1.3.6)^41^ packages.

16S rRNA gene amplicon sequences were analyzed in R Statistical Software (v4.2.1)^37^ with DADA2 (v1.26.0) following the 1.8 Tutorial and Big Data Tutorial^42^. Default parameters for DADA2 were used, except the following: at the 5’ end reads were trimmed 17 bp and truncated at 290 bp (F), while at the 3’ end, reads were trimmed 21 bp and truncated at 200 bp (R). Bacterial ASVs were aligned to the SILVA database (version 138.1)^43,44^ for taxonomic assignment. If ASVs were unassigned at the domain level, they were removed from further analysis. Using phyloseq (v.1.42.0)^45^, all chloroplast, mitochondrial, or plant-assigned ASVs were also removed, leaving 6060 ASVs. Due to varying library sizes, samples were rarefied to 1593 reads per sample and ASVs with fewer than two reads across the entire dataset were removed from further analyses, resulting in 811 unique ASVs across all samples. Using phyloseq, the Simpson and Shannon diversity indices of the microbiomes for each sample were calculated and significant differences were tested for using two separate two-way Scheirer– Ray–Hare tests: seed treatment × developmental stage and seed treatment × chamber type. P-values were adjusted for multiple comparisons using the false discovery rate (FDR) correction. Beta diversity and dispersion were calculated and tested between treatment groups with PERMANOVA and ANOVA, respectively, using both vegan (v.2.6-4)^39^ and pairwiseAdonis (v.0.4.1)^46^. Using indicspecies (v.1.8.0)^47^, ASVs that can be used as indicators for eelgrass development at Stages 1, 3, and 6 were identified. The microbiome package (v.1.20.0)^48^ was used to calculate the relative abundances for all ASVs, and indicator ASVs were visualized with pheatmap (v.1.0.12)^49^. To compare which ASVs were significantly different in relative abundance between seed treatment groups at Stage 6, DESeq2 (v.1.38.3)^50^ was used on non-rarefied data (final non-zero ASVs were 2358). To assess microbiome stability between developmental stages, Spearman’s rank correlation was calculated between Stage 1 and Stage 6 mean abundances for each of the DESeq2-flagged ASVs, analyzed separately for each seed treatment group. The phylogenetic tree of the significant ASVs from the DESeq2 analysis was constructed using Mafft (v.7)^51^ and FastTree (v2.2.0)^52^ and visualized with ggtree (v.3.6.2)^53^.

## Results and Discussion

Of the 1,017 seeds that were individually monitored, 187 seeds germinated (18%), which is within the range of observed germination success at similar salinities under laboratory conditions^28^. After sampling for microbiome composition at Stage 1 (n=12), the remainder of the seeds were placed in either the EcoFAB (n=45), petri dish (n=45), or test tube (n=48) chambers and allowed to grow until the plant developed the first true leaf (Stage 3) or when the cotyledon senesces and abscises, roots are formed, and at least two true leaves have appeared (Stage 6; **Fig. 1**). Non-viable plants (**Table S1**) were excluded from the analysis as mortality was not in the scope of this study.

**Figure 1.**
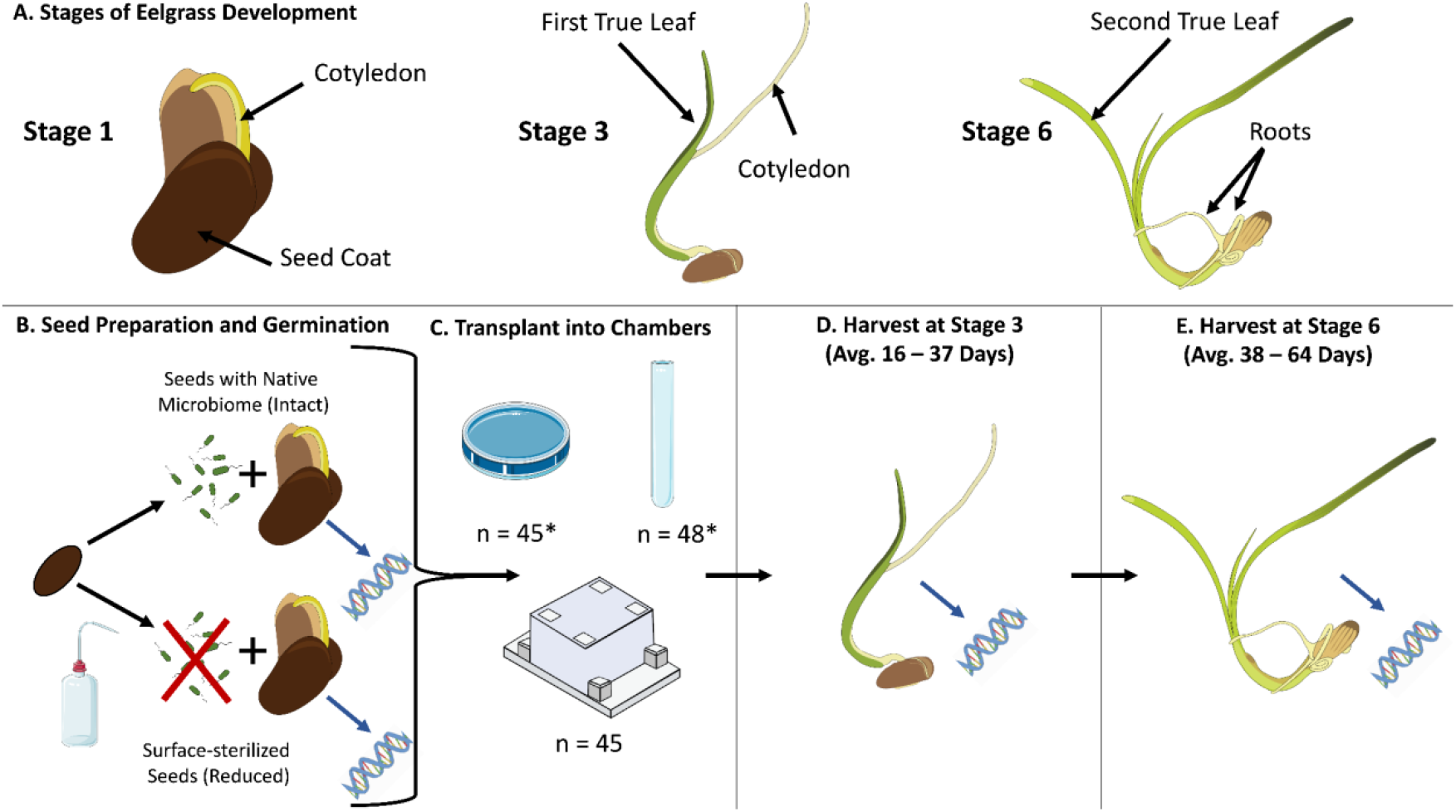
Experimental design and sample breakdown. (**A**) Stages of eelgrass development were based on descriptions by Xu *et. al* 2016^28^. Stages of interest were Stage 1 (initial germination, before introduction to chambers), Stage 3 (when the first true leaf had developed, and Stage 6 (when the cotyledon had senesced and abscised, roots had formed, and the second true leaf had appeared). (**B**) Seeds were prepared to have either reduced or intact microbiomes prior to germination. (**C**) Following germination, plants were randomly selected to be destructively harvested at Stage 1 (indicated by blue arrows) or introduced to EcoFAB 2.0 devices, petri dishes, or test tubes. (**D**) Plants were harvested at either Stage 3 or at (**E**) Stage 6. Each plant was photographed, and morphometric measurements were taken using ImageJ. Following imaging, DNA was extracted from whole plant samples for 16S rRNA gene amplicon sequencing. * Indicates chambers were shaken for 30 minutes per day.

### Eelgrass can be grown reliably in a controlled laboratory setting

We first examined whether seed surface sterilization (reduced microbiome) or light exposure would impact seed germination and seedling development. Seeds with a reduced microbiome had higher germination success (23.2%) in comparison to those with an intact microbiome (13.5%; GLM, df = 1013, p = 0.0019). Germination success was not affected by light conditions (GLM, df = 1013, p = 0.4942) or the interaction between light and seed treatment (GLM, df = 1013, p = 0.5973). There was no effect of the seed surface treatment on *Z. marina* morphological parameters (**Fig. 2**), including days taken to reach each developmental Stages 3 and 6 (Scheirer–Ray–Hare test, p > 0.05), cotyledon length at Stage 3 (Scheirer–Ray– Hare test, p > 0.05), leaf length at either Stage 3 (LMM, p >0.05) or Stage 6 (ANOVA, df = 36, p > 0.05), and root length at Stage 6 (Scheirer–Ray–Hare test, p > 0.05). Our findings indicate that while eelgrass with reduced microbiomes exhibit increased germination, this does not affect plant development; therefore, sterilization is a promising approach for manipulating the seed surface microbiome.

**Figure 2.**
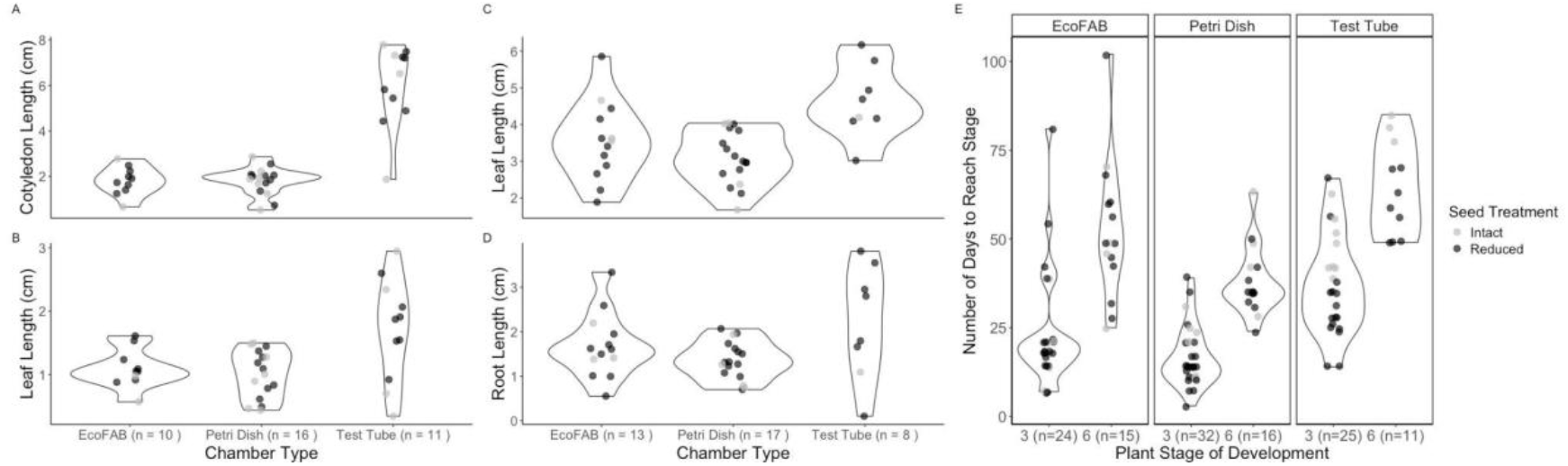
Chamber type, not seed treatment, affects plant morphology and development. Violin plots showing **(A)** cotyledon length (including hypocotyl), **(B)** leaf length at Stage 3, **(C)** leaf length at Stage 6, **(D)** root length at Stage 6, and **(E)** days until development Stages 3 and 6 are reached for eelgrass grown in either EcoFABs, petri dishes, or test tubes.

We used EcoFAB 2.0 devices, petri dishes, and test tubes to determine how different chamber types affected plant development from germination to juvenile seedling. We found that chamber type significantly altered plant morphology (**Fig. 2**). Specifically, Stage 3 cotyledon lengths were significantly longer in test tubes compared to EcoFABs and petri dishes (Scheirer– Ray–Hare test p = 0.00010; Dunn’s Test p < 0.05). A possible reason for this length difference between chambers is the behavior of the axial hypocotyl, which elongates when in an oxygen-reduced environment, such as when buried within sediment^54,55^. Although both test tubes and petri dishes were gently shaken for 30 minutes each day, test tubes may have had lower oxygen concentrations than the petri dish or EcoFAB chamber due to their elongated shape and deeper water column. For leaf length, Stage 3 leaves were not statistically different between chambers (LMM, p >0.05), but plants held in test tubes had significantly longer leaves compared to petri dishes and EcoFABs at Stage 6 (ANOVA, df = 35, p = 0.00174; Dunn’s test p < 0.05). Root length at Stage 6 did not differ between chamber types (Scheirer–Ray–Hare test, p >0.05).

The rate of plant development differed among chamber types (Scheirer–Ray–Hare test p < 0.001 and 0.00028 for Stage 3 and Stage 6, respectively). In test tubes, plants reached Stage 3 after an average of 37 days and progressed to Stage 6 in 64 days. In EcoFABs, development to Stage 3 occurred more quickly, with 23 days, and Stage 6 was reached in 52 days. Petri dishes showed the fastest development, with Stage 3 reached at 16 days and Stage 6 in 38 days. One potential explanation is that with increased cotyledon length, energy reserves for the seed are diminished^54,55^, which could explain the delay in development between stages in test tubes. Together, these findings suggest that test tubes are not ideal chambers for examining seedling development due to the likely stress-induced morphologies being exhibited and the large variability in plant development rates among individuals.

EcoFABs and petri dishes produce seedlings with similar growth and morphologies, but EcoFABs offer significant benefits for seedling-microbiome studies over petri dishes. For example, in petri dishes, plants grow horizontally, and we observed that seedlings began to curve at the sheath as their leaves developed (**Fig. 3**). Plant tropisms such as curving result from multiple plant behaviors, including hormone production^56^, which in turn can affect plant-microbiome interactions^57^. These plant responses add uncertainty in interpreting microbial results and limit the representativeness of findings when extrapolated to what is occurring in the field. Furthermore, in petri dishes, the leaves and roots share the same compartment, eliminating the ability to separate root and leaf environments. This single compartment makes it challenging to sample or manipulate the phyllosphere and rhizosphere independently and allows the two to interact in ways that would not happen in nature. In contrast, the EcoFAB dual-compartment structure separates the plant’s leaves and roots. The upper and lower chambers can be accessed separately through ports without disturbing the plant. They also allow vertical growth, giving seedlings the space to develop more naturally. In the context of microcosm experiments, these features make the EcoFAB a more versatile chamber to use in comparison to Petri dishes.

**Figure 3.**
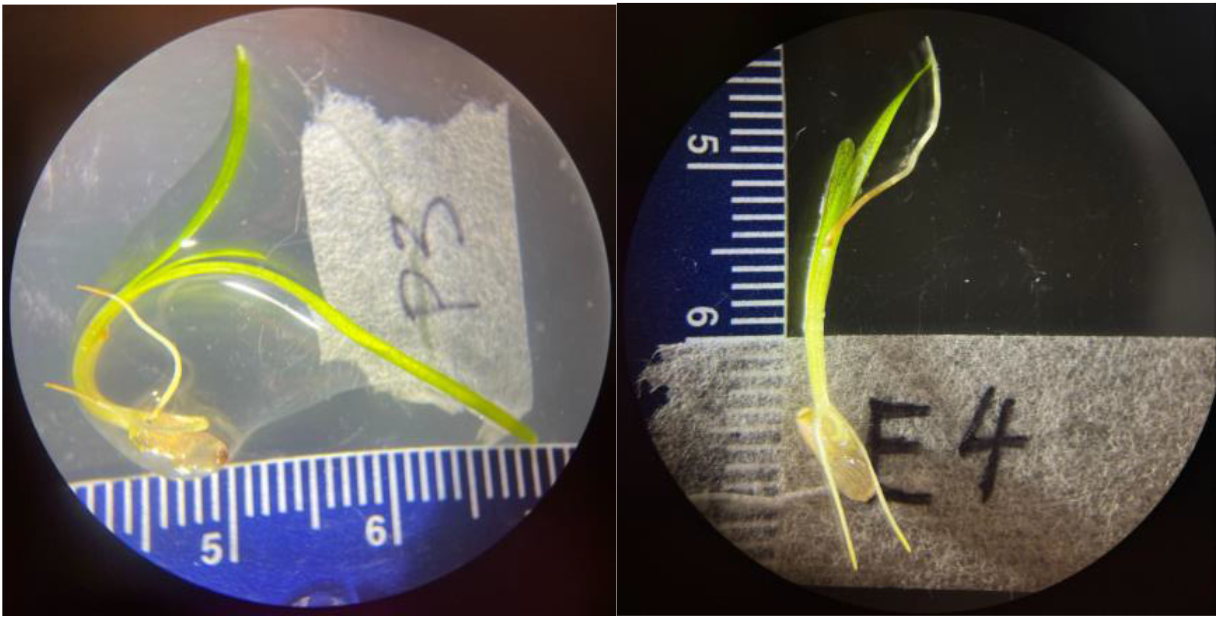
Eelgrass grown in petri dishes (left) displayed greater curvature at the sheath as leaves developed compared to those grown in EcoFABs (right). Representative dissecting microscope images (10x magnification) of eelgrass seedlings (Stage 6) in either petri dishes (left) or EcoFAB 2.0 devices (right). Scale bar = 0.1 mm.

### Eelgrass microbiome assembly can be manipulated with sterilization techniques and has long-term trajectory effects

To determine whether surface sterilization (reduced microbiome) of *Z. marina* seeds altered microbiome assembly trajectories throughout plant development, we analyzed overall microbial richness and evenness (alpha diversity) as well as community composition (beta diversity; **Table 1**). Although we included chamber effects in our analyses, given plant morphological differences between chamber types (**Fig. 2**), our primary focus here centers on seed treatment affecting the microbiome assembly as the plant grows, regardless of chamber type, as this is most relevant for future EcoFAB-based studies. Chamber type and developmental stages of eelgrass did not impact the overall alpha diversity (Scheirer–Ray–Hare test; p > 0.05, FDR corrected), but seeds with reduced microbiomes had significantly lower alpha diversity (Scheirer–Ray–Hare test: Simpson diversity p = 0.004, Shannon diversity p = 0.0001, FDR corrected; **Fig. 4A-B**). For beta diversity, we observed significant differences between eelgrass grown from seeds with reduced microbiomes versus those with intact microbiomes (**Fig. 4C-D**; **Table 1**; PERMANOVA, p < 0.05). All variables except the seed treatment and stage interaction effect were also statistically significant. Additionally, seedlings from seeds with reduced microbiomes showed greater microbiome dispersion across individuals than those from seeds with intact microbiomes (**Fig. 4C**; PERMDISP, p = 0.0039).

**Table 1.** Seed treatment, chamber type, and plant developmental stage significantly affect the microbiome composition. Permutational multivariate analysis of variance (PERMANOVA) results based on Bray-Curtis dissimilarity of Hellinger-transformed ASV count data, testing the effects of seed treatment, chamber type, and plant developmental stage on microbiome community composition. The model used was ∼ Seed treatment* Chamber + Seed treatment* Chamber + Stage + Chamber * Stage, including main effects and significant interactions. Seed treatment, chamber type, and development stage were significant factors, along with the seed treatment × chamber type and chamber type × stage interactions. Results are based on 10,000 permutations.

|  | Df | Sum of Squares | R<sup>2</sup> | F | Pr(>F) |
| --- | --- | --- | --- | --- | --- |
| **Seed Treatment** | 1 | 2.6784 | 0.09312 | 8.0058 | **9.999e-05** |
| **Chamber Type** | 3 | 2.0401 | 0.07093 | 2.0326 | **0.000200** |
| **Stage** | 1 | 0.5383 | 0.01871 | 1.6088 | **0.034997** |
| **Seed Treatment:Chamber Type** | 3 | 1.5070 | 0.05239 | 1.5015 | **0.006599** |
| **Seed Treatment:Stage** | 1 | 0.2919 | 0.01015 | 0.8725 | 0.633437 |
| **Chamber Type:Stage** | 2 | 0.9654 | 0.03356 | 1.4428 | **0.031597** |
| **Residual** | 62 | 20.7428 | 0.72114 |  |  |
| **Total** | 73 | 28.7639 | 1 |  |  |

**Figure 4.**
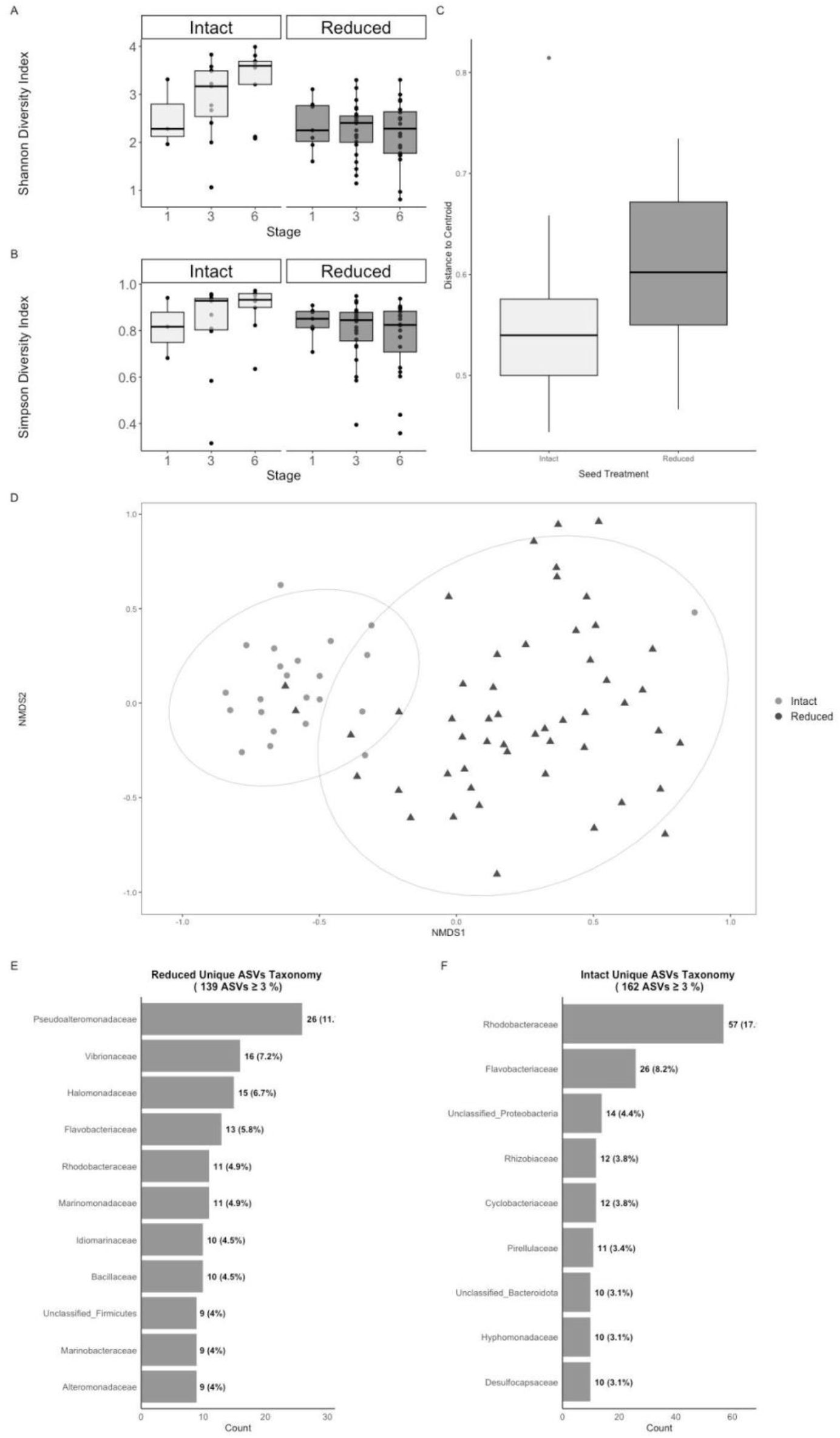
Eelgrass from surface-sterilized seeds (Reduced) have a lower alpha diversity and a higher beta-dispersion in their assembly than those from seeds with native microbiomes (Intact). **(A)** Boxplots of Simpson and **(B)** Shannon alpha diversity of eelgrass microbiomes across plant Stages 1, 3, and 6 within seed treatment (Reduced and Intact). **(C)** Dispersion between plants with an intact versus reduced microbiome. **(D)** NMDS plot using Bray-Curtis dissimilarity of Hellinger-transformed ASV count data. (**E and F**) Horizontal bar charts displaying the family-level taxonomic composition of ASVs unique to each seed treatment group across developmental stages. Numbers in parentheses indicate the percentage of unique ASVs within each family. Families only representing more than 3% of the unique ASVs are represented.

The observed reductions in alpha diversity and increases in inter-sample variation following sterilization are comparable to what has been previously found with terrestrial plant seed sterilization protocols in plant-microbe studies, which remove native bacterial epiphytes ^58,59^ to allow targeted microbial additions^60,61^. In our system, while the identity of surviving taxa after sterilization appeared to be stochastic, we expect that these remnant microorganisms would be displaced by taxa introduced at high density in future inoculation experiments.

To understand which taxa were driving the diversity differences between eelgrass from treated and untreated seeds, we first examined ASVs only present on plants with either reduced (223 ASVs) or intact microbiomes (319 ASVs). For families representing >3% of these unique ASVs, only Rhodobacteraceae and Flavobacteriaceae were observed in both treatments (**Fig. 4D and 4E**). Plants with intact microbiomes had higher proportions of Rhodobacteraceae (17.9%) and Flavobacteriaceae (8.2%) among their unique ASVs, while these families comprised only 5.8% and 4.9% of unique ASVs in reduced microbiomes, respectively. Both families are known to be associated with older eelgrass leaves, but are also found on many marine surfaces, which would explain their greater representation in intact seed microbiomes^62^. In addition, eelgrass that had intact microbiomes had treatment-specific ASVs belonging to families typically associated with the rhizosphere and roots (i.e., Rhizobiaceae, Cyclobacteriaceae, and Desulfocapsaceae) or leaves (i.e., Hyphomonadaceae and Pirellulaceae) of the mother plant^17,63,64^. This pattern reflects the plant material that seeds would have been exposed to during initial release in the mesh bags.

To further explore the trajectory of microbiome assembly between treatment groups as they near establishment, we analyzed plants at Stage 6 when both leaves and roots were fully developed. We used DESeq2 to identify individual ASVs that were significantly different in relative abundance between plants with reduced microbiomes versus those with intact microbiomes (log2 fold-change > 2; p-adjusted < 0.05). Out of 2358 ASVs, 38 had significantly higher relative abundances for eelgrass in the reduced microbiome treatment versus the intact group, while 105 ASVs had higher relative abundances for plants in the intact microbiome treatment (**Fig. 5; Table S2**).

**Figure 5.**
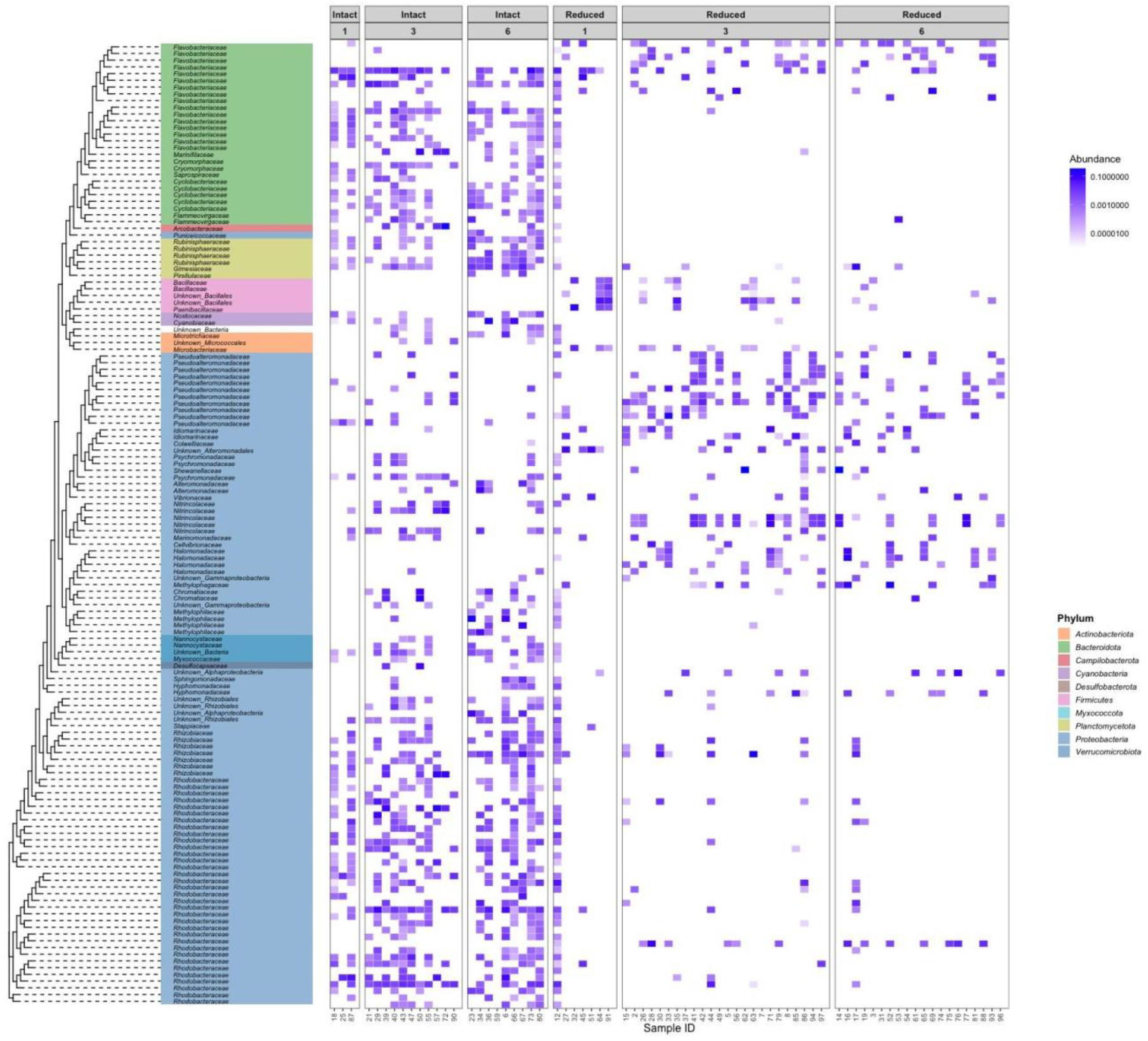
Eelgrass from native microbiome seeds (Intact) maintain higher relative abundances of taxa from Stage 1 onward, unlike those from surface-sterilized seeds (Reduced). Heat map of the relative abundance (log10 transformed) of the 143 significantly different ASVs between Reduced and Intact plants at Stage 6, regardless of chamber type. Relative abundances are plotted across all developmental stages.

ASVs enriched in eelgrass with reduced microbiomes included members from the family Halomonadaceae (log2 fold change >24), which harbors many taxa with plant growth-promoting properties^65^. For example, members of the Halomonadaceae contribute to plant salt tolerance, as these bacteria reduce the negative impacts of saline stress through the production of exopolysaccharides and other compounds, such as ectoine, that enhance seed germination under high saline conditions^66–69 70–72^. Due to being enriched in eelgrass from seeds with reduced microbiomes and taxa reported as endophytes in other plants^73–75^, we suggest that Halomonadaceae are important endophytic members of *Z. marina* during its early development.

Among the 105 ASVs enriched in the intact microbiome treatment, Rhizobiaceae emerged as a particularly notable family, with members showing substantial enrichment (log2 fold change 8.9-45) relative to the reduced group. The presence of Rhizobiaceae, along with the previously identified Rhodobacteraceae and Flavobacteriaceae, reflects the epiphytic community derived from mother plant material. Rhizobiaceae, which include Rhizobia, are known for nitrogen fixation and facilitate ammonium mobilization in the *Z. marina* rhizosphere^76^. In general, there are higher amounts of nitrogen-fixing bacteria found in seagrass meadows compared to unvegetated sediment^77^, with juvenile plants having higher relative abundances compared to adults^16^. Therefore, bacteria with these capabilities may be important for seedling establishment. How epiphytes explicitly establish on the plant post-germination is largely unknown (but see War *et al*. 2023 for a full review), but their high relative abundances highlight their potential importance^78^.

We observed a pattern throughout the plant development for the 143 ASVs flagged by our DESeq2 analysis: The microbiomes on plants developing from untreated seeds established rapidly and remained stable throughout the experiment, at least with respect to the dominant taxa by Stage 6. However, when microbiomes were reduced on seed surfaces, the resulting microbiome changed across plant development stages (**Fig. 5**). To further explore the stability of the microbiome between plant stages, we calculated Spearman’s rank correlation between Stage 1 and Stage 6 for the average abundance of each ASV for plants with reduced microbiomes versus intact microbiomes to see if ASVs that had high abundances at Stage 1 remain high at Stage 6. For intact microbiomes, Spearman’s rho was 0.4391 (p = 4.12 e-08), suggesting that there was a moderate to strong ASV abundance consistency. Plants with reduced microbiomes, however, had a Spearman’s rho of 0.09 (p =0.264), suggesting no correlation between stages. Overall, our results support that we can alter the *Z. marina* microbiome with seed-surface sterilization by decreasing the microbial diversity of the seed and altering the assembly dynamics of the microbiome throughout seed development.

### Eelgrass development causes shifts in the microbiome composition

Beyond seed treatment effects, we sought to understand how host filtering shapes microbiome assembly during plant development. To investigate the trajectory of the microbiome during development, we restricted the analysis to surface-sterilized seeds grown in EcoFABs due to the interaction effect of stage and chamber type (p=0.0146; **Table 2**). By focusing on plants from seeds with reduced microbiomes, we can determine how seed derived endophytes assemble with limited impact of epiphytes derived from the environment. In addition, delving into this portion of the microbiome gives insight to potential bacterial members that are vertically transmitted from the mother to seeds. 16S rRNA gene amplicon sequencing of plants from sterilized seeds revealed that microbiome composition significantly differed between Stages 1, 3, and 6 (PERMANOVA p=0.0043, **Fig. 6A**). We identified 26 ASVs in our indicator species analysis that are significantly associated with the three *Z. marina* developmental stages (**Fig. 6B**; **Table S3**). Of note, members of the Halomonadaceae family were found in Stages 3 and 6. Halomonadaceae are expected to be increasing in abundance with plant development, likely providing support for osmotic regulation, stress signaling, and hormonal balance as seen for other plant hosts^79^. Additionally, members of the Rhodobacteraceae family were found in Stage 6. Rhodobacteraceae are commonly associated as marine surface colonizers^23^, but some species have been identified as endophytes for salt-tolerant plants *Salicornia europaea* and *Populus euphratica* Oliv., suggesting these bacteria may play specialized roles in salt tolerance during eelgrass seedling development. Of the 12 indicator ASVs for Stage 1, 10 belonged to Bacillaceae and Paenibacillaceae. These same two families are dominant families (39 and 35.6%, respectively) in tomato seeds, and some species exhibit strong antifungal properties, suggesting their importance in seed protection^80^. Furthermore, both families have shown to have capabilities to inhibit oomycetes^81,82^, which are common Zostera seed pathogens^83,84^. This stage-specific recruitment of functionally diverse bacterial families, including protective and stress-tolerant, demonstrates how marine plants assemble specialized microbial partnerships to address the multifaceted challenges of early development. Seed endophytes can drive a large portion of microbiome assembly in terrestrial plants, which is speculated to result from priority effects and vertical transmission^10,14,85–87^; here, we found that similar processes can occur in seagrass seeds. This effect was particularly pronounced for seeds that had their initial microbiome reduced. Even if these form only a small component of the overall community in untreated seeds in nature, the putative key role these taxa play in stage-specific challenges may still affect the overall health of early life stages.

**Table 2.** Permutational multivariate analysis of variance (PERMANOVA) results based on Bray-Curtis dissimilarity of Hellinger-transformed ASV count data, testing the effects of chamber and stage on plants grown from seeds with reduced microbiomes. The model used was ∼ Chamber Type * Stage. Results are based on 10,000 permutations.

|  | Df | Sum of Squares | R<sup>2</sup> | F | Pr(>F) |
| --- | --- | --- | --- | --- | --- |
| **Chamber Type** | 3 | 1.9184 | 0.10083 | 1.8042 | **0.0007** |
| **Stage** | 1 | 0.3984 | 0.02094 | 1.1241 | 0.2817 |
| **Chamber Type:Stage** | 2 | 1.1137 | 0.05854 | 1.5711 | **0.0146** |
| **Residual** | 44 | 15.5950 | 0.81969 |  |  |
| **Total** | 50 | 19.0255 | 1 |  |  |

**Figure 6.**
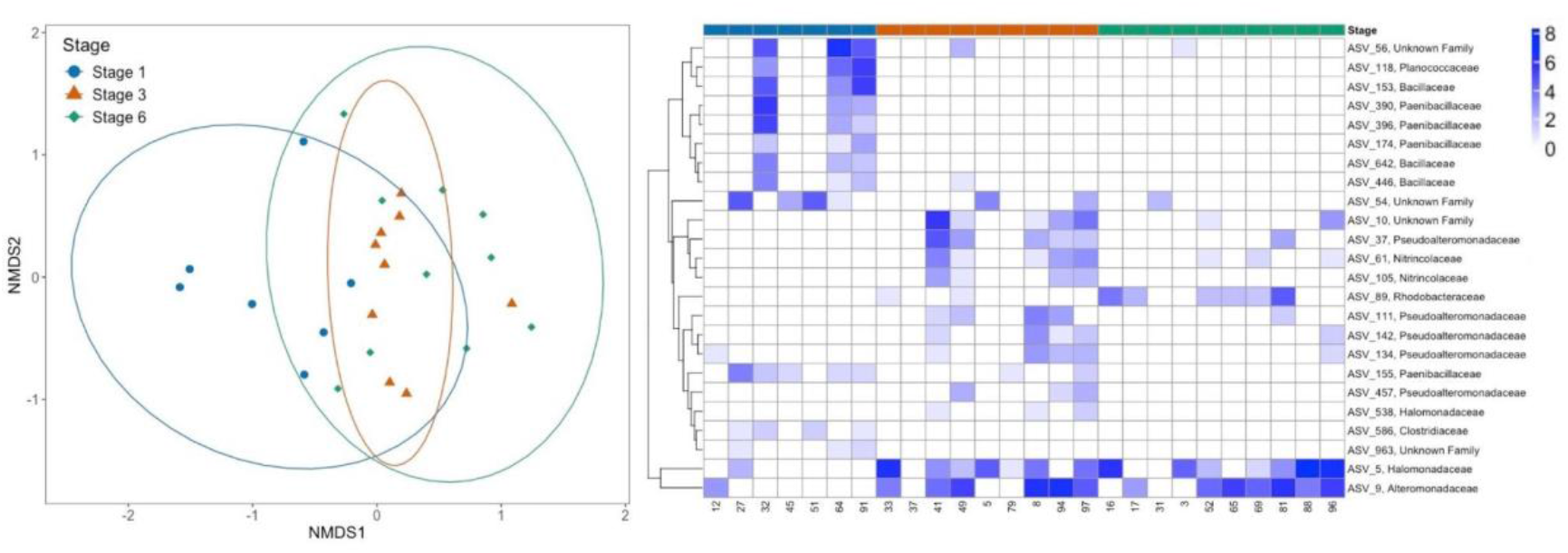
Microbiome composition shifts with host development. **(A)** NMDS plot based on Bray-Curtis dissimilarity of Hellinger-transformed ASV count data showing separation of microbiome across plant stages grown from reduced microbiome seeds. **(B)** Heat map showing the relative abundance (log(1+x) transformed) of the indicator species determined for each stage of the plant.

## Conclusion

In this study, we successfully grew eelgrass in a controlled laboratory setting and were able to manipulate and detect the microbiome shifts throughout early developmental life stages using the EcoFAB 2.0. Establishing a standardized methodology for microbiome reduction in seagrass seedlings without harming host morphology is essential. This work lays the foundation for future microcosm studies focusing on introducing beneficial microbes as synthetic communities to *Z. marina* seedlings. Based on our results, we recommend adding synthetic communities to the seed coat rather than later in plant development to establish a long-term presence of the desired microbes. However, further work is needed to see if this pattern holds when other biological factors are included. For example, microbes from the sediment could out-compete the synthetic community for functional niches or host filtering at later plant stages could further alter the microbiome composition (e.g. reproductive stage)^13,86,88^.

Identifying both putative endophytes and epiphytes at different stages of plant growth is essential for experimentally testing the role of beneficial bacteria in plants ^89–94^. In our work, we show that eelgrass development had a stronger influence on endophytic than epiphytic community composition, and that the epiphytic community had many overlapping taxa in microbiome studies of adult seagrass, which suggests vertical transmission^95^. We further highlight families with potentially important roles: endophytic Halomondaceae for salt tolerance, epiphytic Rhizobiaceae for N acquisition, and endophytic Paenibacillaceae for pathogen protection. Further exploration of microbiome dynamics among taxa such as these is needed to determine which interactions have synergistic or antagonistic effects on the host^96^.

Beyond identifying and testing bacterial candidates for desired host functions, we can leverage the EcoFAB 2.0 to explore plant metabolisms by manipulating abiotic and biotic conditions across plant compartments. While this study focuses on the microbiome of the entire plant (leaf and root tissue together), the EcoFAB 2.0’s dual chamber setup also enables separate quantification of seagrass root and shoot microbiome shifts. This compartmentalized approach provides opportunities to examine plant-microbe interactions in both the phyllosphere and rhizosphere, ultimately leading to a more comprehensive understanding of how specific microbial members contribute to overall seagrass health. For example, the lower chamber could be manipulated to contain varying nutrients not available in the upper chamber, such as sulfides, that are known to affect seagrass meadow health^97,98^. The lower chamber could also house sediment-derived bacteria, mimicking what the seed would be exposed to as it disperses from the mother plant, allowing for investigation of how eelgrass microbiomes assemble when native seed members are competing with environmental sources of new species. Together, these capabilities position the EcoFAB 2.0 as an instrumental tool for defining critical seagrass-microbe interactions, ultimately advancing our understanding needed to inform and optimize restoration efforts in these coastal ecosystems.

## Acknowledgements

We’d like to thank Diane Brache-Smith (UC Merced) for her contributions to Figure 1. This work was supported by funding from NSF (NSF BioOCE #2311577) and the Gordon and Betty Moore Foundation (Marine Model Systems #9359). The work (proposal: CSP New Investigator Proposal 509268) conducted by the U.S. Department of Energy Joint Genome Institute (https://ror.org/04xm1d337), a DOE Office of Science User Facility, is supported by the Office of Science of the U.S. Department of Energy operated under Contract No. DE-AC02-05CH11231.

## Competing Interest

P.F.A. and T.R.N. are inventors on patent US11510376B2 held by University of California that covers Ecosystem device for determining plant-microbe interactions. T.R.N. and B.P.B. are scientific co-founder and hold equity in Datacodec with prior approval from LBNL.

## Supplemental Data

**[Table S1. Seed Assignment and Data Collection By Date](https://ucdavis.box.com/s/gzrki5a1hd1yg6h3s65oo8jrldi5ek67)**

**[Table S2. DESeq2 Analysis Output](https://drive.google.com/file/d/1lD1KSONyvsDfbV3ijiqRMBBQ-B4m-9t4/view?usp=sharing)**

**[Table S3. Indicator Species Analysis Output](https://ucdavis.box.com/s/ca61rozulgmat3diu9a7i3tu4stcpoin)**

## Notes

### Summary of Updates

Figure 1 has been added, and minor revisions for the conclusion.

